# Cluster mean-field theory accurately predicts statistical properties of large-scale DNA methylation patterns

**DOI:** 10.1101/2021.09.03.458837

**Authors:** Lyndsay Kerr, Duncan Sproul, Ramon Grima

## Abstract

The accurate establishment and maintenance of DNA methylation patterns is vital for mammalian development and disruption to these processes causes human disease. Our understanding of DNA methylation mechanisms has been facilitated by mathematical modelling, particularly stochastic simulations. Mega-base scale variation in DNA methylation patterns is observed in development, cancer and ageing and the mechanisms generating these patterns are little understood. However, the computational cost of stochastic simulations prevents them from modelling such large genomic regions. Here we test the utility of three different mean-field models to predict large-scale DNA methylation patterns. By comparison to stochastic simulations, we show that a cluster mean-field model accurately predicts the statistical properties of steady-state DNA methylation patterns, including the mean and variance of methylation levels calculated across a system of CpG sites, as well as the covariance and correlation of methylation levels between neighbouring sites. We also demonstrate that a cluster mean-field model can be used within an approximate Bayesian computation framework to accurately infer model parameters from data. As mean-field models can be solved numerically in a few seconds, our work demonstrates their utility for understanding the processes underpinning large-scale DNA methylation patterns.

## 1 Introduction

DNA methylation is a repressive epigenetic mark [1] which is primarily found on the cytosines of CpG dinucleotides in mammals. Double-stranded CpG dyads can be unmethylated or methylated on both strands (*u* and *m* respectively) or methylated on only one strand (hemimethylated, *h*). DNA methylation is largely erased from the genome during early mammalian development [2]. It is then re-established by the *de novo* DNA methyltransferases DNMT3A and DNMT3B [3] resulting in a landscape where 70-80% of CpGs are methylated in most human cells [4]. Regulatory elements such as promoters and enhancers often remain methylation free [1]. During DNA replication, the nascent strand is synthesised with unmethylated cytosines and methylation patterns are copied by the maintenance methyltransferase, DNMT1 [5]. Failure to maintain DNA methylation at a locus results in passive DNA demethylation. Methylation can also be removed actively through transient modification by Ten Eleven Translocation (TET) enzymes and subsequent DNA repair [6].

Waves of demethylation and remethylation take place during early development and the generation of germline cells [2]. Changes in DNA methylation patterns also occur during development and cellular differentiation, resulting in cell type specific methylation patterns [7]. The correct establishment of DNA methylation patterns is vital for normal development. Mutations in DNMTs cause Mendelian disorders in humans [8, 9, 10] and mice knockouts die before or shortly after birth [3, 5]. Widespread alterations in DNA methylation patterns occur in cancer and ageing [11, 12], but the significance of these changes is unclear. It has been observed that globally hypomethylated mice expressing a single hypomorphic DNMT1 allele develop cancer, suggesting that altered DNA methylation can cause cancer [13]. However, the mechanisms underpinning DNA methylation changes remain unclear preventing the robust delineation of their role in development and disease.

Mathematical models are powerful tools for understanding complex biological processes, including DNA methylation. The importance of interactions between CpGs in maintaining DNA methylation patterns was first postulated through modelling [14]. Specifically, the authors modelled collaborative interactions where CpGs within a region of the genome can influence the state of other CpGs, e.g. through enzyme recruitment. Models including such collaborativity were subsequently found to explain experimental measurements of methylation maintenance *in vitro* and *in vivo* more closely than those that did not include it [15, 16]. A recent study also suggests that collaborativity mediated by neighbour-guided error correction through DNMT1 is important for maintaining DNA methylation [17]. Deterministic models, non-spatial stochastic models and spatial stochastic models have all been used to describe DNA methylation [18]. Deterministic models are based on rate equations while stochastic models are based on Fokker-Planck equations or chemical master equations (CMEs). CMEs are ideal because they take into account the inherent discreteness of molecular fluctuations [19] which is well known to play an important role in cellular dynamics [20]. The CME of simple non-spatial stochastic models can be solved exactly in closed-form [21], but this is often not possible for spatial stochastic models. Rather in this case, stochastic simulations are used to model the individual reaction processes described by the CME. Various types of stochastic models have been used to describe collaborative methylation systems (see for example [22, 23, 24, 25, 26]). To date, such mathematical models have been applied to understand methylation patterns on a kilo-base scale. However, mega-base scale alterations to DNA methylation patterns occur in development, cancer and ageing [27]. Existing models rely on simulations that are too computationally expensive to run for such large genomic regions.

Here we test the idea that large-scale steady-state methylation patterns can be modelled in a tractable manner using mean-field (MF) models. By comparison to synthetic data generated from stochastic simulations, we demonstrate that a type of cluster mean-field model can predict the statistical properties of large-scale methylation patterns. In Section 2 we introduce a nearest-neighbour collaborative model for DNA methylation and describe the process used to simulate data from this model. We describe the three MF models we test in Section 3. In Section 4 we compare the ability of each MF model to predict statistics associated with methylation patterns resulting from the simulations. We find that a type of cluster mean-field model provides excellent predictions and demonstrate that this model can be used within an Approximate Bayesian Computation (ABC) framework to infer parameters underpinning large-scale methylation systems. Finally, in Section 5, we discuss the implications of our findings.

## 2 Nearest-neighbour collaborative model

### 2.1 Model set-up

We consider the reaction system in Fig. 1, where some reactions are non-collaborative (involving only the “target” CpG, whose methylation state changes during the reaction), while others are collaborative (involving both a target CpG and a “mediator” CpG). The role of the mediator is to encourage the reaction to occur, e.g. via the recruitment of methylase or demethylase enzymes. This system, and reduced versions, have previously been used to examine small-scale methylation patterns [14, 28, 29].

**Figure 1:**
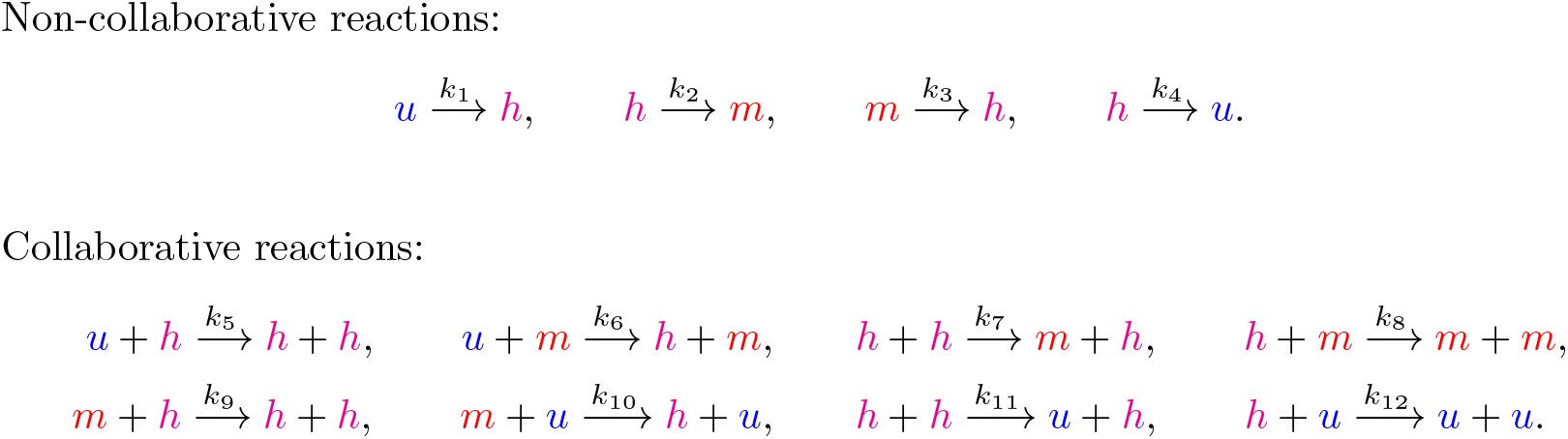
System of reactions under consideration. Here *u, h* and *m* represent unmethylated, hemimethylated and methylated CpGs, respectively. Non-collaborative reactions involve only one CpG, while collaborative reactions involve two CpGs. For each collaborative reaction, the second reactant (the mediator—see text) recruits an enzyme that changes the methylation state of the first reactant (the target). For example, the reaction *u* + *h* → *h* + *h* involves a hemimethylated CpG at one site recruiting an enzyme which changes the state of a CpG at another site from unmethylated to hemimethylated. Reaction rates are *k*_*i*_, *i* = {1, …, 12}.

We make the following assumptions:

i. *CpGs can only influence the methylation state of their nearest neighbours; see Fig. 2*. While both experimental and modelling studies have demonstrated the importance of CpGs being influenced by surrounding CpGs [14, 15], the extent and range of such influence is unknown. We therefore consider only interactions between nearest neighbours.
ii. *There are no direct transitions between the unmethylated and methylated states*. We justify this with the observation that methylase and demethylase enzymes act on single DNA strands [6, 30]. This implies that hemimethylation is a necessary transition state between unmethylated and fully methylated CpGs.
iii. *The system has reached a steady state*. Here, we assume that there are no long-term effects of DNA replication on methylation patterns. This assumption is supported by the observation that the DNA methylation patterns of cycling and arrested cells are similar [31].

**Figure 2:**
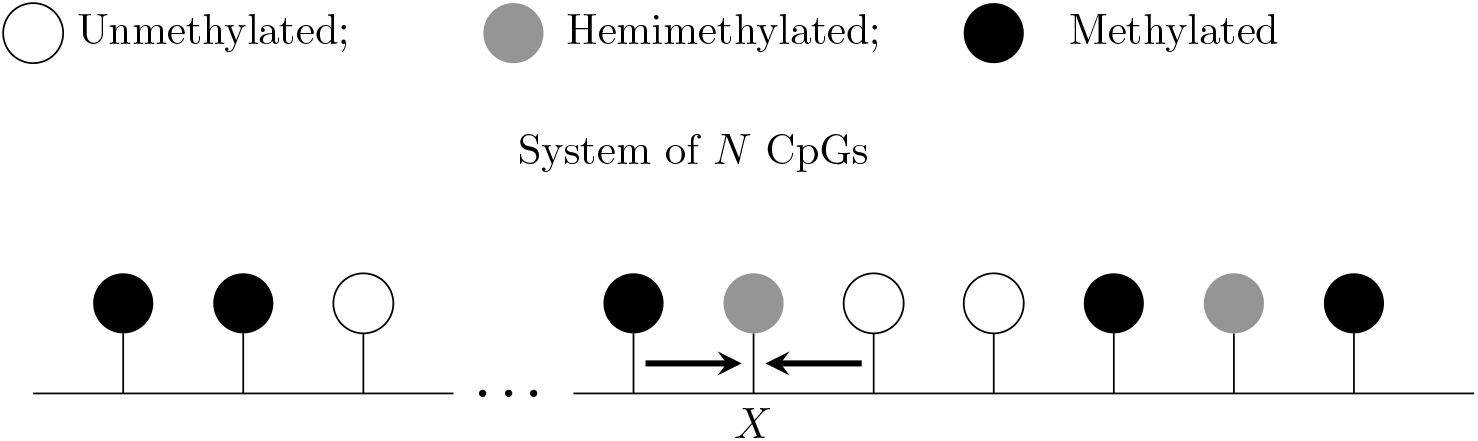
Collaborative interactions that can influence a target *X* under the nearest-neighbour collaborative model. Individual CpGs are represented by “lollipops”, with their colour indicating their methylation status (white: unmethylated, grey: hemimethylated and black: methylated). Collaborative methylation and demethylation reactions can only occur between neighbouring CpGs (potential influences on CpG *X* are indicated by arrows). There is no upper bound on *N* ∈ ℕ, allowing large-scale methylation patterns to be considered.

We also assume that the rates *k*_*i*_, *i* = {1, 2, …, 12} are of the form given in Table 1, where *x* measures the strength of collaborativity between CpGs (*x* < 1 indicates that non-collaborative reactions dominate, while *x* > 1 indicates that collaborative reactions dominate). The parameter *y* measures the strength of methylation vs. demethylation (*y* < 1 corresponds to demethylation dominating and *y* > 1 corresponds to methylation dominating). The parameter *a* scales the reaction rates.

**Table 1:** Reaction rates for the model in Fig. 1.

|  | Demethylation | Methylation |
| --- | --- | --- |
| Non-Collaborative | $k_3 = k_4 = a$ | $k_1 = k_2 = ay$ |
| Collaborative | $k_9 = k_{10} = k_{11} = k_{12} = ax$ | $k_5 = k_6 = k_7 = k_8 = axy$ |

### 2.2 Simulations of nearest-neighbour collaborative system

In the simple case of two CpGs, the system can be in six possible states: *mm, uu, hh, um, hm* and *uh*. For such a small system, all possible transitions between states (via reactions in Fig. 1) can be identified and the evolution of the system can be described exactly by six mathematical equations, one for each state. However, for large systems, it is infeasible to identify all possible states and transitions between states meaning that equations describing the exact evolution of the system can not be formulated. While stochastic simulations are computationally expensive, they are the ground truth of the nearest-neighbour collaborative system to which we compare our MF models and so here we describe the process underlying these simulations.

We focus on the steady-state case so that *u, h* and *m* levels fluctuate around some fixed steady-state values. For a system of *N* CpGs, we simulate the nearest-neighbour collaborative system using Gillespie’s algorithm [32]—see Fig. 3 for an illustration of how this algorithm simulates a 4-CpG system. A sample taken at any time point will contain *N* methylation states. For each parameter set, we therefore sample the system at a total of *T* = 10^6^*/N* time points after steady state has been reached to obtain a dataset of 10^6^ steady-state methylation states.

**Figure 3:**
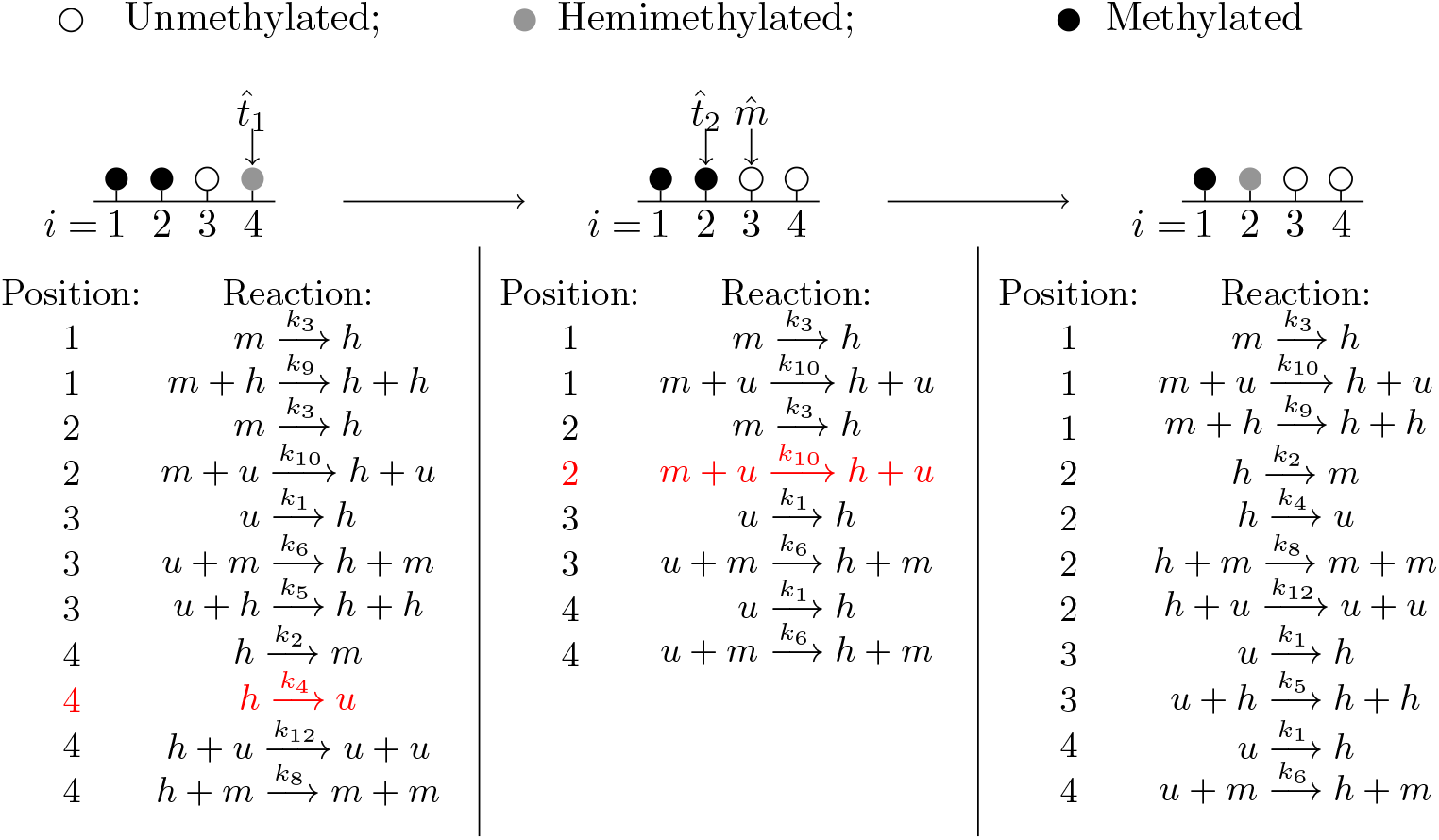
Stochastic simulations of the nearest-neighbour collaborative system. For simplicity, a 4-CpG system is considered here. We fix *a, x* and *y* and impose periodic boundary conditions so that the first and final CpG can interact. All potential reactions are first identified, and the Gillespie algorithm is used to choose one of these reactions, and a time for it to occur. The system is updated to account for the reaction occurring and the process is repeated to generate dynamical behaviour. In the example shown, we start with the system on the left. All possible reactions are listed and a 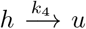 reaction is chosen to occur at target position 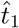 The system and the list of potential reactions are then updated (middle). Subsequently, another reaction, 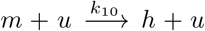, is chosen to occur at target position, 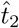, with CpG 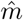 acting as mediator. The system and the list of possible reactions is again updated (right). This process is repeated until the system reaches steady state.

### 2.3 Analysis of simulated data

To facilitate our analysis, we define a sequence *u*^*t*^, *t* ∈ {1, …, *T*}, where

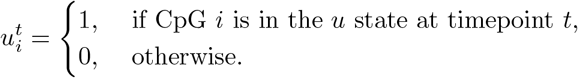

We define *h*^*t*^ and *m*^*t*^ similarly; see Fig.4. We also define *z*^*t*^ via

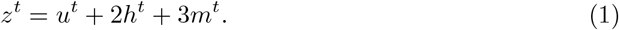

**Figure 4:**
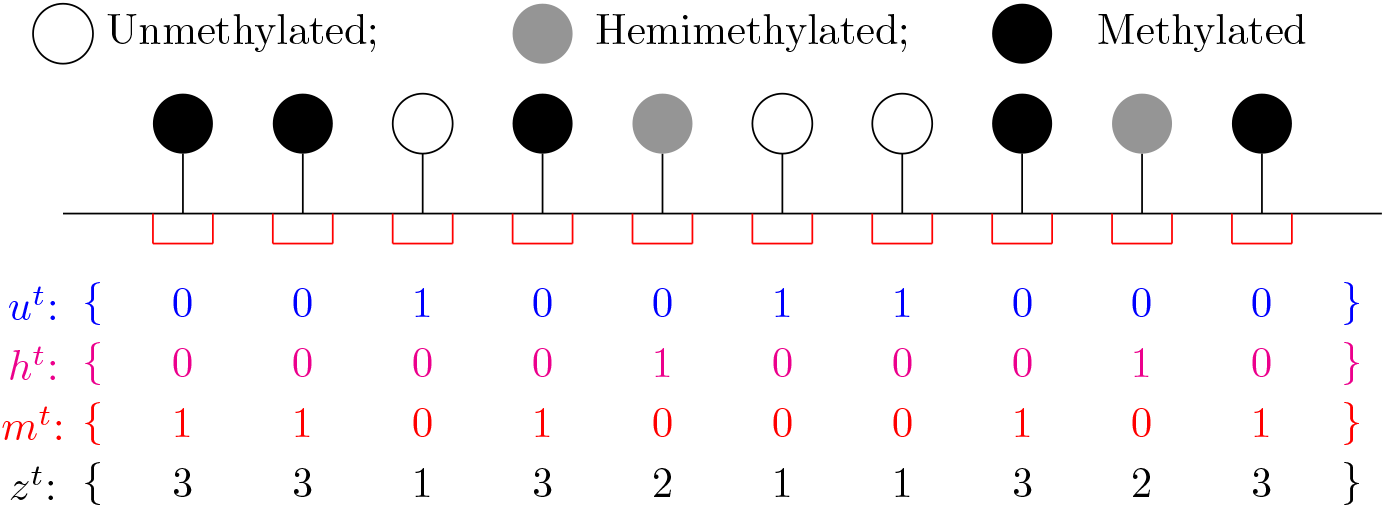
Construction of the sequences *u*^*t*^, *h*^*t*^, *m*^*t*^ and *z*^*t*^. For simplicity, only ten CpGs are shown for a single time point, *t*. A vector *u*^*t*^ is created, where 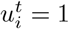 if CpG *i* is unmethylated at time *t* and 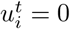 otherwise. Vectors *h*^*t*^ and *m*^*t*^ are constructed similarly. Finally, a vector *z*^*t*^ is created via *z*^*t*^ = *u*^*t*^ + 2*h*^*t*^ + 3*m*^*t*^.

The mean *u, h* and *m* levels, *µ*_*us*_, *µ*_*hs*_, *µ*_*ms*_, are obtained via

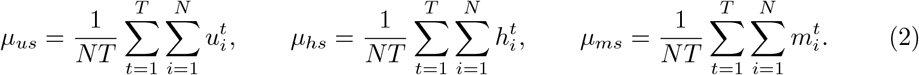

For each *t*, we also calculate the mean and variance over (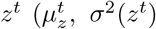, respectively) and average over all *t* ∈ {1, …, *T*} to obtain an overall mean and variance, *µ*_*z*_ and σ^2^(*z*), which are given by

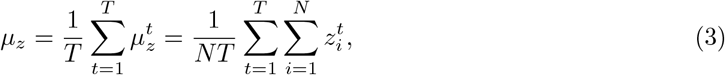

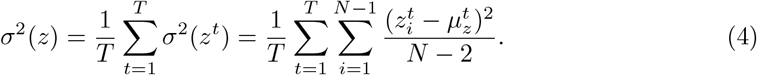

We then define sequences *v*^*t*^ and *w*^*t*^ by

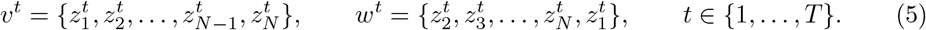

For each *t* ∈ {1, …, *T*}, the sequence *v*^*t*^ is identical to *z*^*t*^, while *w*^*t*^ is a shifted version of *z*^*t*^ (i.e. 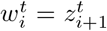 for *i* = {1, … *N* − 1} and 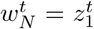). For each *i* = {1, …, *N*}, comparing 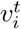 and 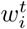 provides information regarding the methylation state of two neighbouring CpGs at time *t* ∈ {1, …, *T*}. We calculate the covariance between *v*^*t*^ and *w*^*t*^ for each *t*, Covar(*v*^*t*^, *w*^*t*^), averaging over *t* ∈ {1, …, *T*} to obtain an overall covariance between neighbouring sites

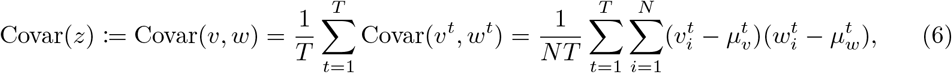

where 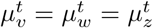. Finally, the correlation between neighbouring sites, *ρ*(*z*) := *ρ*(*v, w*), is calculated via

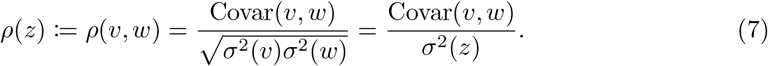

## 3 Mean-field models for DNA methylation maintenance

For large CpG systems, the CME describing the nearest-neighbour collaborative model cannot be easily solved and stochastic simulations are computationally expensive. In contrast, it is often the case that models simplified using the MF approximation can be computationally solved in a time-efficient manner and we aim to test whether they accurately approximate the nearest-neighbour collaborative model for DNA methylation. To this end, we construct three MF models (see Sections 3.1, 3.2, 3.3). These models consider an infinite system of CpGs and so, by design, their ability to accurately describe a genomic region increases with the size of the region. In these models, nearest-neighbour interactions are approximated by considering the mean state of the system. In the first model, nearest-neighbour interactions are entirely approximated by considering the probability that two states are adjacent (one-site MF model). The second model describes distinct pairs of CpGs (distinct pairs MF model). Interactions occurring within a pair are directly accounted for and other nearest-neighbour interactions are approximated by considering the probability that two paired states are adjacent. In the third model we consider overlapping pairs of CpGs (overlapping pairs MF model). Interactions occurring within a pair are again directly accounted for, but now other nearest-neighbour interactions are approximated by considering the probability that two paired states overlap. The remainder of this section is devoted to mathematical descriptions of these models.

### 3.1 One-site mean-field model

We define the proportion of sites in the *u, h, m* states to be the mean *u, h, m* levels, *µ*_*u*_, *µ*_*h*_, *µ*_*m*_, respectively. Here we construct a one-site MF model, where changes in the system are influenced by *µ*_*u*_, *µ*_*h*_, *µ*_*m*_, rather than nearest-neighbour interactions; see Fig. 5.

**Figure 5:**
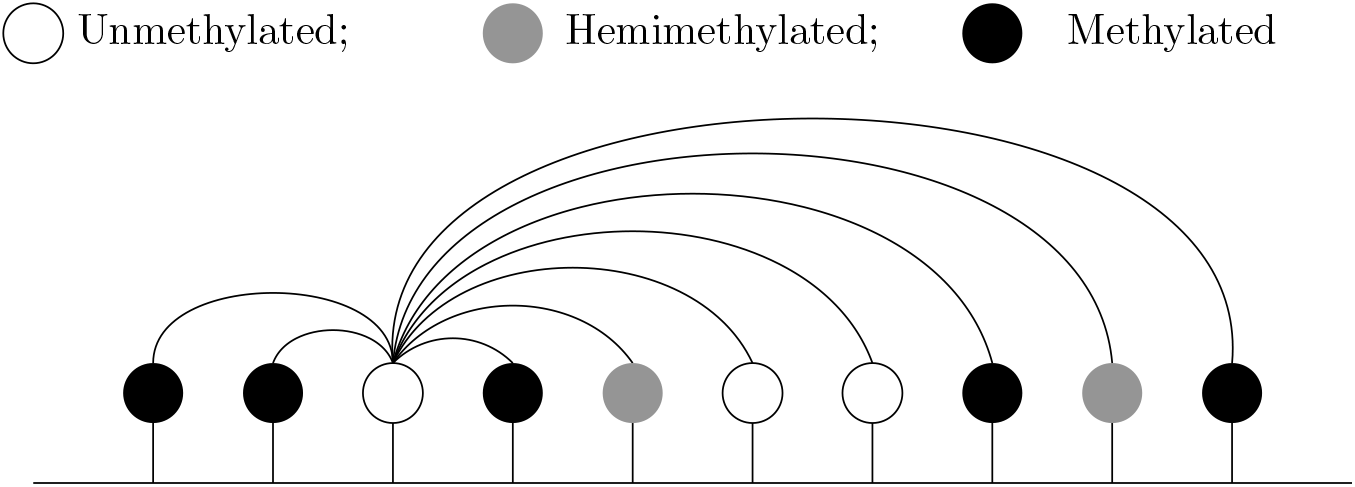
Schematic of the one-site MF model. CpGs are influenced by the mean of the system rather than nearest-neighbour interactions.

Consider the reaction 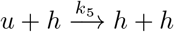 in Fig. 1. Since the *h* mediator is unchanged by the reaction, we can write this as an effective first-order reaction 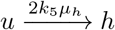, where the *h* mediator is absorbed into the effective reaction rate by making it proportional to *µ*_*h*_. The factor of two accounts for the *h* mediator being on either side of the *u* target. Similarly, 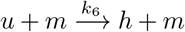 can be written as 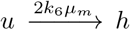. Thus the 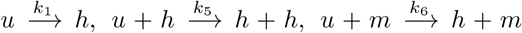 reactions in Fig. 1 can be written as a single effective first-order reaction 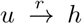, with *r* = *k*_1_ + 2*k*_5_*µ*_*h*_ + 2*k*_6_*µ*_*m*_. We thus write the system in Fig. 1 as the effective first-order system

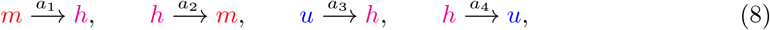

where

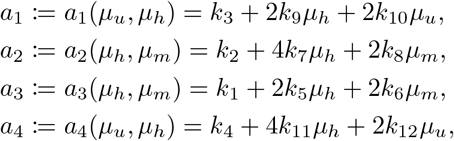

and *µ*_*u*_ + *µ*_*h*_ + *µ*_*m*_ = 1. The *k*_7_ and *k*_11_ terms have an additional factor of two since their associated reactions involve two *h* reactants, and either of these can change state during the reaction.

Let *L*_*u*_, *L*_*h*_, *L*_*m*_ be the “level” (proportion) of *u, h, m* at a single CpG, respectively. A CpG can only be in one state at any time and so

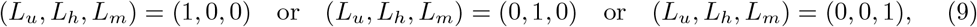

for each CpG. Using (8) we construct a CME describing the probability that a site is in the *u, h* or *m* state and from this we can obtain moment equations (see Appendix C of Ref [33]), for the statistics of *L*_*u*_, *L*_*h*_, *L*_*m*_. The mean values are *µ*_*u*_ = ⟨*L*_*u*_⟩, *µ*_*h*_ = ⟨*L*_*h*_⟩, *µ*_*m*_ = ⟨*L*_*m*_⟩, where the angled brackets denote the expected value. Due to the conservation law *L*_*m*_ = 1 − *L*_*u*_ − *L*_*h*_, we need only consider equations for *u* and *h*. The means are described by the equations,

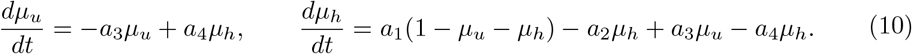

Setting

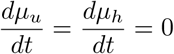

leads to implicit equations for the steady-state means,

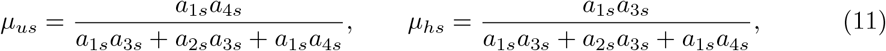

where *a*_1*s*_ = *a*_1_(*µ*_*us*_, *µ*_*hs*_), *a*_2*s*_ = *a*_2_(*µ*_*hs*_, 1 − *µ*_*us*_ − *µ*_*hs*_), *a*_3*s*_ = *a*_3_(*µ*_*hs*_, 1 − *µ*_*us*_ − *µ*_*hs*_), *a*_4*s*_ = *a*_4_(*µ*_*us*_, *µ*_*hs*_). Since Eq. (11) is independent of *a*, the means depend only on *x* and *y*. For fixed parameters, we can solve Eq. (11) numerically to obtain values for *µ*_*us*_ and *µ*_*hs*_.

The second moment equations are given by

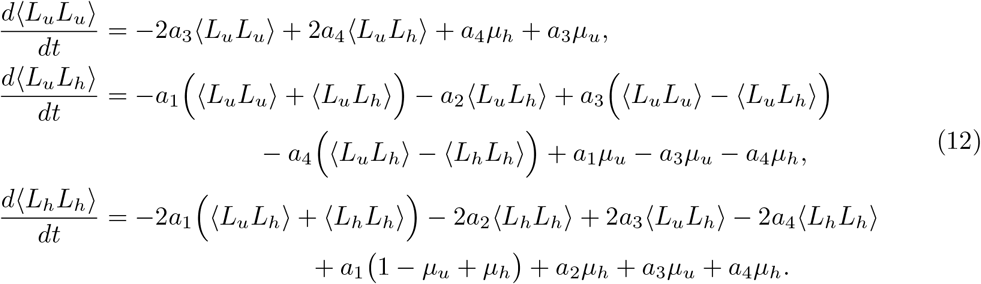

From Eq. (9) we expect *L*_*u*_*L*_*u*_ = *L*_*u*_, *L*_*h*_*L*_*h*_ = *L*_*h*_ and *L*_*u*_*L*_*h*_ = 0 at any CpG. Solving Eq. (12) in steady state leads to

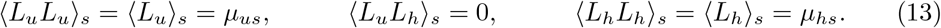

Variances, σ^2^(*L*_*us*_) and σ^2^(*L*_*hs*_), can then be obtained via

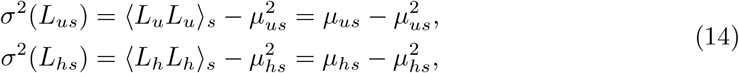

along with the covariance Covar(*L*_*us*_, *L*_*hs*_), which is given by

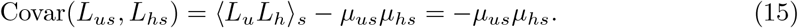

The mean, variance and covariances associated with the *m* state can now be obtained using

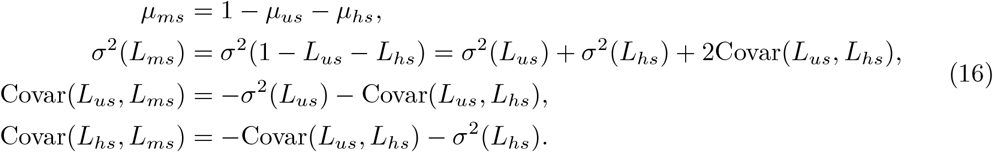

We calculate the steady-state mean and variance, *µ*_*z*_ and σ^2^(*z*), associated with the variable *z* = *L*_*u*_ + 2*L*_*h*_ + 3*L*_*m*_ via

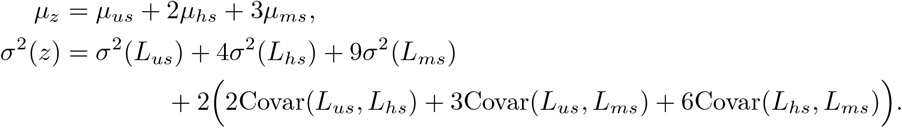

Note that the superscript *t* was only used in Section 2.2 to differentiate between samples at different time points. Here we simply have a single *z*. Since no spatial information is obtained from the one-site MF model, the covariance and correlation between neighbouring sites cannot be extracted.

### 3.2 Distinct pairs mean-field model

We next construct a two-site MF model, where we consider “clusters” of two adjacent CpGs. Such cluster MF models have been successfully used to study vehicular traffic and driven-diffusive gas models [34, 35]. We define the mean level (proportion) of pairs in the six possible states,

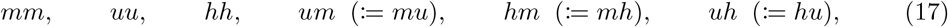

to be *µ*_1_, *µ*_2_, *µ*_3_, *µ*_4_, *µ*_5_, *µ*_6_, respectively. Here *µ*_4_ is the proportion of pairs containing *u* and *m*, irrespective of order. Similarly, *µ*_5_ is the proportion of pairs containing *h* and *m*, and *µ*_6_ is the proportion of pairs containing *u* and *h*, irrespective of order.

In the distinct pairs MF model (DPMF model), CpGs within a pair are allowed to interact directly, preserving some nearest-neighbour interactions. The influence of the nearest-neighbour CpGs flanking the pair is then approximated by considering the probabilities that an adjacent pair is in each of the six possible states; see Fig. 6. Here, each CpG belongs to only one pair and each pair of sites is a single reactant. As with the one-site model, we consider an effective first-order reaction system, given by

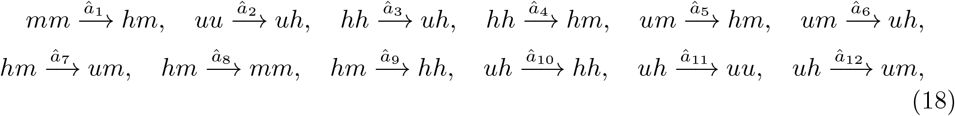

**Figure 6:**
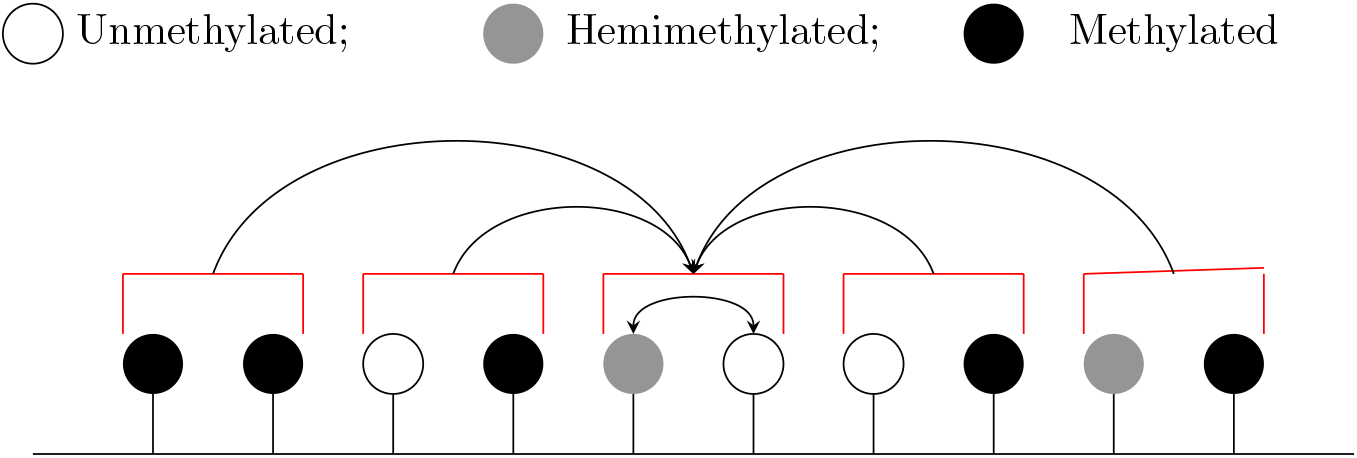
Schematic of the distinct pairs MF model. The two CpGs within a pair can interact directly with each other and the pair is also influenced by the mean state of pairs in the system. In the figure, the *uh* pair can change state due to interactions between the *u* and *h* within the pair, and due to the mean state of pairs in the system.

where the effective rates are given by

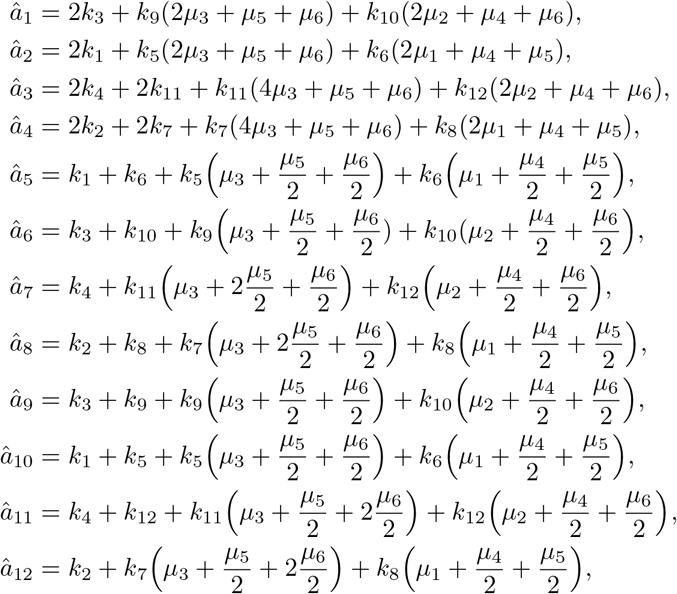

and 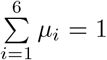. We describe the construction of *â*_1_ in Appendix A. Essentially, *â*_1_—*â*_12_ are constructed by considering all possible ways that each reaction in (18) can occur via a reaction from Fig. 1 taking place. Such reactions can occur within the reactant pair or can take place between a site within the pair and a site from an adjacent pair. Terms associated with interactions between two *hh*, two *hm* or two *uh* pairs have an additional factor of two since either pair can change state during the reaction. While we can, in principle, calculate the distribution of pairs in (18) [36], we restrict our attention to obtaining moments of the system.

Let *L*_1_, *L*_2_, *L*_3_, *L*_4_, *L*_5_, *L*_6_ be the level (proportion) of each of the paired states at a single pair of CpGs. At any time, a pair can be in only one state and so at a single pair we have

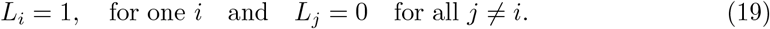

The *u, h, m* levels within a pair of CpGs, 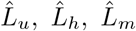, are then given by

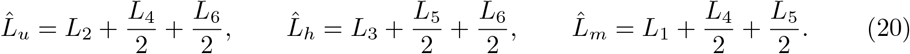

Using the CME for the system (18) we construct first and second moment equations for *L*_*i*_, *i* = {1, …, 6}; see Appendix C. The first moment equations describe *µ*_*i*_ = ⟨*L*_*i*_⟩, the mean values of *L*_*i*_ for *i* = {1, …, 6}. For fixed parameters, solving these equations numerically in steady state gives the steady-state means, *µ*_*is*_, *i* = {1, …, 6}. Note that these means are independent of *a*.

From the second moment equations, we obtain the steady-state expected values of *L*_*i*_*L*_*j*_, ⟨*L*_*i*_*L*_*j*_⟩_*s*_, for *i, j* ∈ {1, 2, …, 6}. As expected from Eq. (19), ⟨*L*_*i*_*L*_*i*_⟩_*s*_ = ⟨*L*_*i*_⟩_*s*_ = *µ*_*is*_, ⟨*L*_*i*_*L*_*j*_⟩_*s*_ = 0 for all *i* ≠*j, i, j* ∈ {1, 2, …, 6}. Variances and covariances are then obtained using

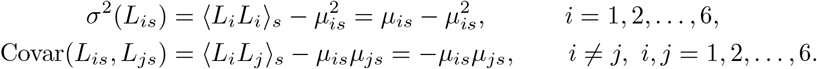

Once again, these are independent of the parameter *a*.

We now have statistics for the paired states in (17). The steady-state means for the *u, h, m* levels in a pair are then given by

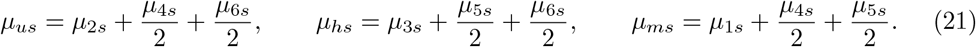

The pair-to-pair variance in *u* level is given by

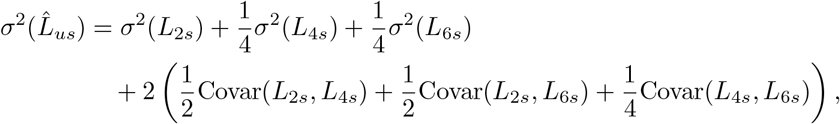

and similarly for the variances associated with *h* and *m*, 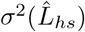 and 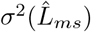. Covariances are given by

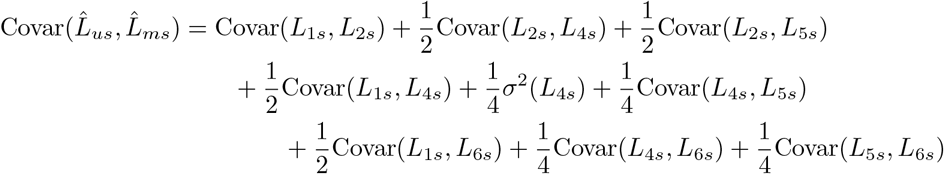

and analogously for 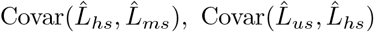.

Note that the statistics obtained so far relate to *u, h, m* levels within a pair of CpGs. The mean level of a state within a pair is the same as the mean level of the state at each site. However, this is not the case for higher moments. For example, 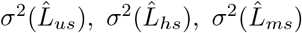 are pair-to-pair variances, rather than site-to-site variances.

We aim to obtain statistics relating to *z* = *L*_*u*_ + 2*L*_*h*_ + 3*L*_*m*_, where *L*_*u*_, *L*_*h*_, *L*_*m*_ are the single-site *u, h, m* levels. We define 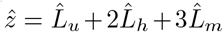, noting that 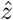 contains information regarding pairs of CpGs. Essentially,

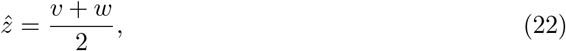

where *v* and *w* are as in Eq. (5) and we again do not require the superscript *t*.

The means of *z, v, w* and 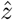 coincide and

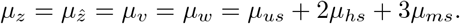

Also, *vw* = *L*_2_ + 2*L*_6_ + 3*L*_4_ + 4*L*_3_ + 6*L*_5_ + 9*L*_1_ and so

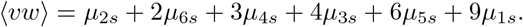

From this we calculate the covariance between neighbouring sites as

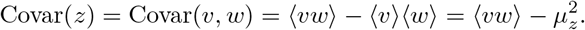

The variance associated with 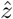 can be calculated via,

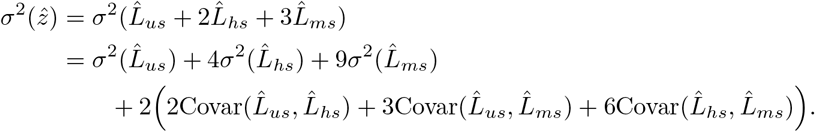

However, 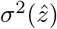 is the pair-to-pair variance. Using σ^2^(*z*) = σ^2^(*v*) = σ^2^(*w*) we obtain

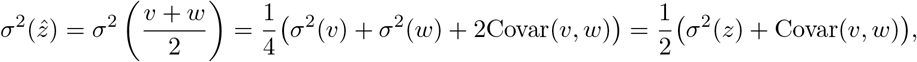

leading to the site-to-site variance,

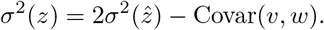

The correlation between neighbouring pairs is obtained via

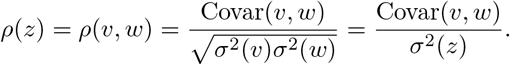

To summarise, the statistics of primary interest from our calculations are: the means *µ*_*us*_, *µ*_*hs*_, *µ*_*ms*_, *µ*_*zs*_, the variance σ^2^(*z*), the covariance Covar(*z*) and the correlation *ρ*(*z*).

### 3.3 Overlapping pairs mean-field model

Similarly to the DPMF model, there are also six possible states for a pair of CpGs in the overlappling pairs MF model (OPMF model), see (17). The OPMF model also incorporates direct interactions within a pair. However, each CpG now belongs to two pairs, one with its left-hand neighbour and one with its right-hand neighbour, leading to a system of overlapping pairs. The influence of CpGs flanking a pair is now approximated by considering the conditional probability that the pair overlaps with another pair of a certain state; see Fig. 7.

**Figure 7:**
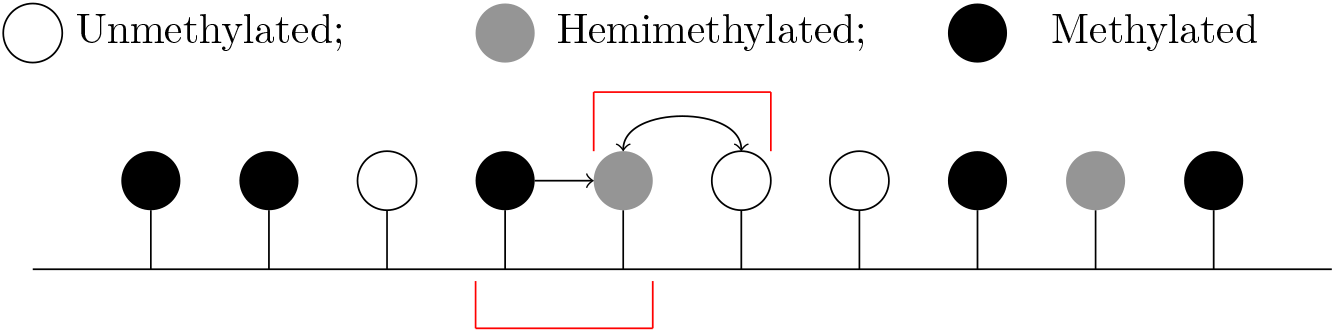
Schematic of the overlapping pairs MF model. A pair of CpGs interact directly with each other and the effect of CpGs flanking the pair is approximated by considering the conditional probabilities that a flanking site is in the *u, h* or *m* state, given the state of its neighbour within the pair.

As before, we consider an effective first-order reaction system, given by

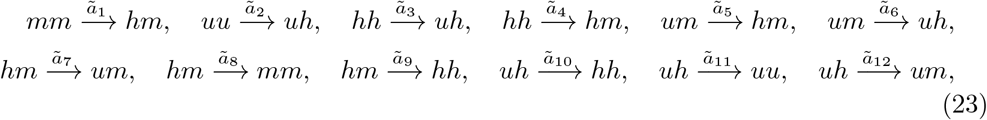

where the effective rates are given by

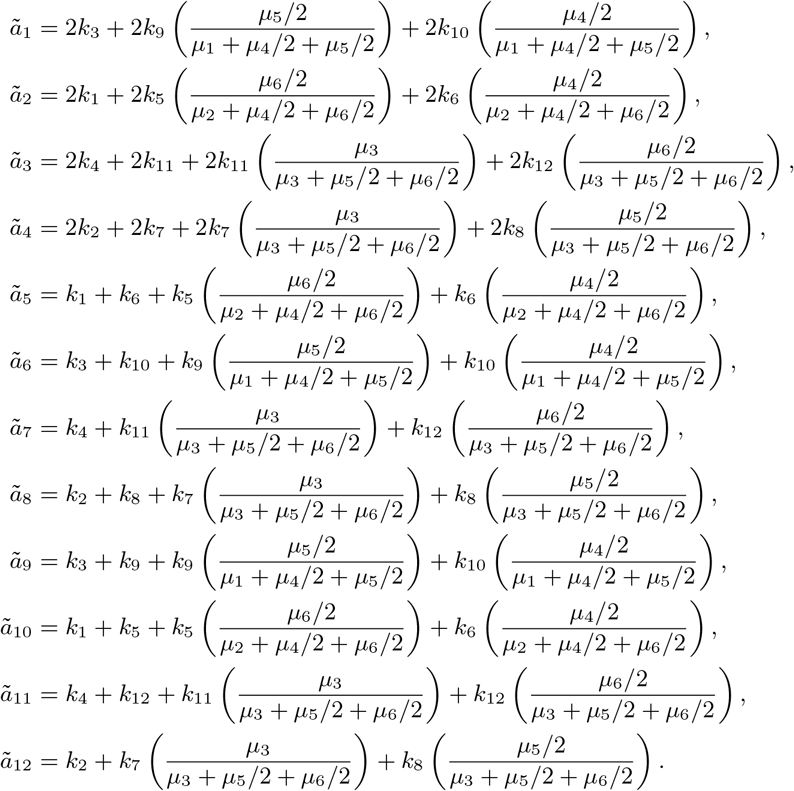

We detail the construction of *â*_1_ in Appendix B.

An identical approach as in Section 3.2 leads to first and second moment equations for the system, see Appendix C, and from these we obtain means, variances and covariances associated with the paired states and with the pair-to-pair *u, h, m* levels. Again, these depend only on *x* and *y*. The statistics associated with *z* are obtained as for the DPMF model, see Section 3.2.

## 4 Model comparison and parameter inference

To test whether MF models are capable of modelling large-scale methylation patterns, we now compare model predictions to synthetic data generated using nearest-neighbour collaborative simulations. The statistical properties obtained from our models are independent of *a* and so we fix *a* = 0.2. Since demethylation dominates when *y* < 1 and methylation dominates when *y* > 1, we hypothesise that a sharp change in the behaviour of the system may occur at *y* = 1. To capture this potential transition for different collaborativity strengths, we consider *x* = {0.1, 1, 5, 50}, *y* = {0.1, 0.2, …, 2}. For every parameter set, we run *n* = 10 stochastic simulations, obtaining ten datasets of 10^6^ CpGs, from which we calculate the statistics of interest. We then calculate the means and standard errors over the ten datasets to obtain overall summary statistics.

In this study, we approximate the nearest-neighbour collaborative system by MF models which consider an infinite system of CpGs. As *x* increases, finite-size effects cause discrepancies between the simulations and models, which can be counteracted by increasing the number of simulated sites. We therefore simulate systems of *N* = 200 sites when *x* = {0.1, 1, 5} and systems of *N* = 500 sites when *x* = 50.

### 4.1 Mean-field models capture steady-state methylation levels

We first compare the mean *u, h, m* levels (*µ*_*us*_, *µ*_*hs*_, *µ*_*ms*_) from the MF models to those from the simulations (Fig. 8). Considering the simulated data first, we observe that *u* and *m* dominate when *y* < 1 or *y* > 1 respectively. *h* is an intermediate state between *u* and *m* and peaks at *y* = 1, where there is also a sharp transition between *u* and *m*-dominant states.

**Figure 8:**
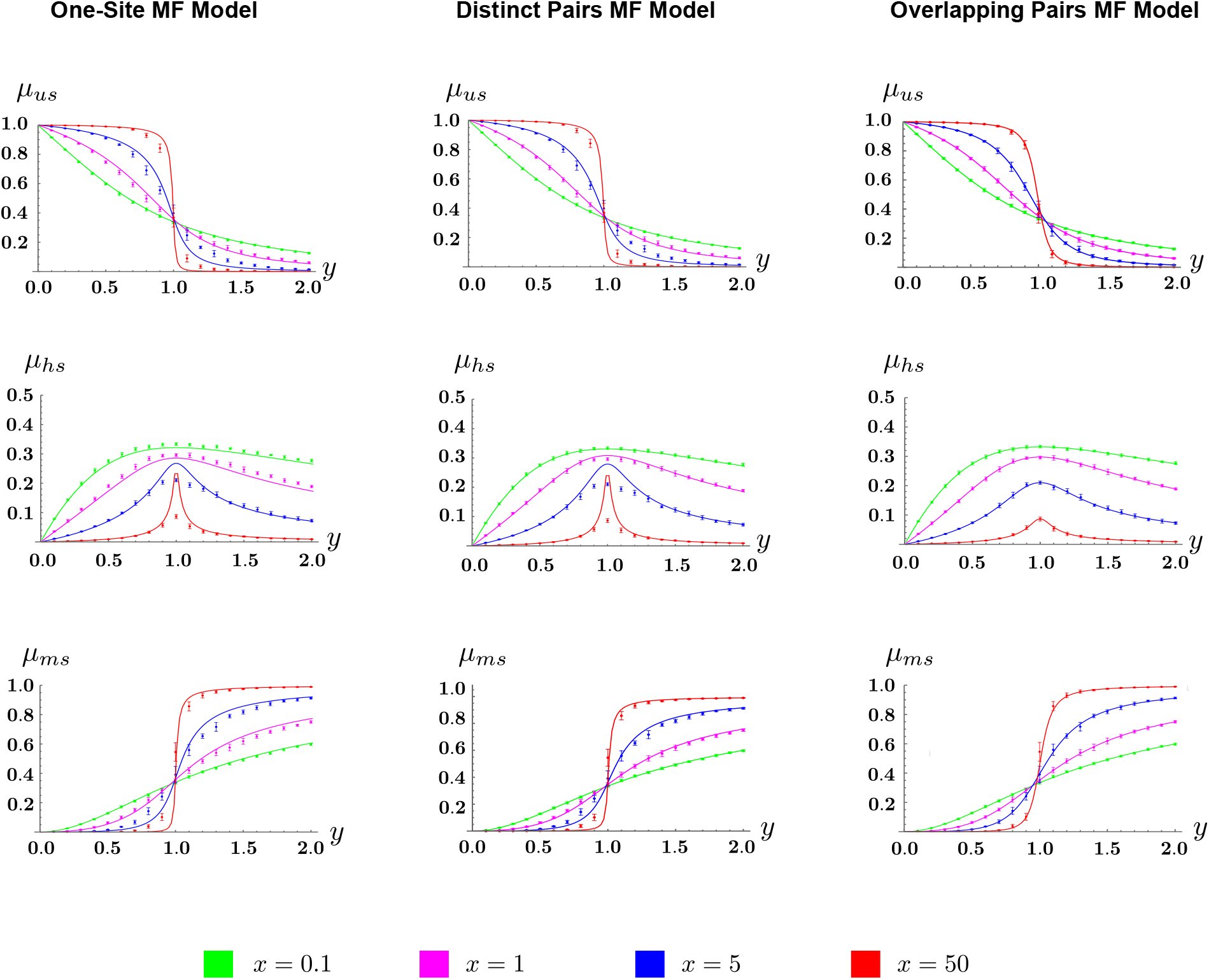
The OPMF model accurately predicts the average behaviour of large-scale methylation levels. The mean *u, h* and *m* levels are plotted against the methylation strength *y* for various values of the collaborativity strength *x* and the different mean-field models (left: one-site MF model, middle: DPMF model and right: OPMF model). Solid lines denote model predictions and points show the mean of *n* = 10 simulations. Error bars indicate standard error for the simulated data.

While all models capture the qualitative behaviour of the means as *x* and *y* are varied, we observe that predictions of methylation levels from the OPMF model are closest to those observed in the simulated data (Fig. 8). All three models predict the mean *u* and *m* levels reasonably well, however only the OPMF model accurately predicts the mean *h* level for large *x* and for *y* close to one. All predictions from the OPMF model are within the error of the simulated data and it successfully captures the transition observed at *y* = 1 for all *x* considered. Conversely, the predictions of the other models deviate from the simulations at the transition point when *x* is large. The one-site MF model deviates to the greatest extent for 87% of the parameter sets. The poorer performance of the one-site model can be seen most clearly in the mean *h* levels when *x* = 1. This suggests that the one-site model has the worst predictive power and we exclude it from further analysis.

### 4.2 The OPMF model accurately predicts associations between neighbouring sites

To test whether MF models can predict associations between neighbouring CpGs we consider *z* = {*z*_1_, *z*_2_, …, *z*_*N*_}, where *z*_*i*_ = {1, 2, 3} if CpG *i* is in the *u, h, m* state, respectively. From *z* we calculate the mean and variance associated with the methylation state, and the covariance and correlation in methylation state between neighbouring sites. In the simulated data (Fig. 9), we again observe a transition in these statistics when the methylation and demethylation strengths are equal (*y* = 1). Our results counterintuitively suggest that neighbouring sites are most correlated here (the peak *ρ*(*z*) occurs when *y* = 1).

**Figure 9:**
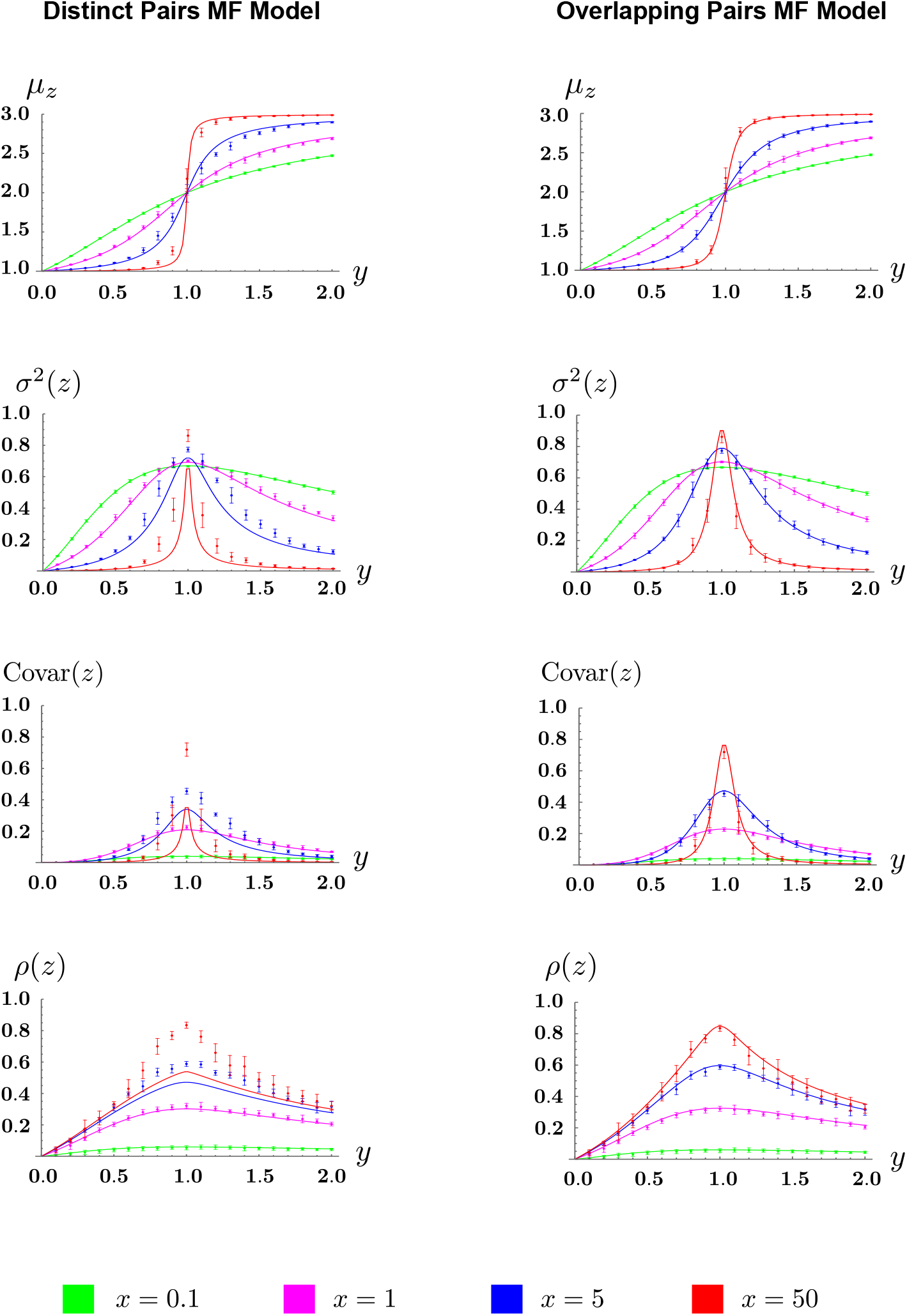
The OPMF model accurately predicts associations between neighbouring CpGs. Predictions of the means (*µ*_*z*_), variances (σ^2^(*z*)), covariances (*Covar*(*z*)) and correlations (*ρ*(*z*)) are plotted against *y* for different values of *x* and for the DPMF model (left-hand panels) and the OPMF model (right-and panels). Solid lines denote model predictions and points show the mean statistics calculated from the simulated data (*n* = 10 in each case). Error bars indicate standard error for the simulated data. Note that, since *z* is determined by *x* and *y*, the statistics plotted here are implicit functions of *x* and *y*.

To gain insight into this observation, we examine the patterns that evolve for *x* = 50 (Fig. 10). When *y* is small, large *u* clusters form and we intuitively expect neighbouring sites to be highly correlated. However, these large clusters are interspersed with infrequent, isolated occurrences of *h* and *m* which have low correlations with their neighbours. Moreover, ten *m* (or *h*) sites appearing in isolation will result in smaller *u* clusters than the ten sites appearing as a single cluster. The overall effect is to reduce the correlation when *y* is small. A similar rationale explains the low correlation when *y* is large. *u* and *m* cluster sizes are most similar when methylation and demethylation are equally strong, resulting in *u* and *m* sites correlating equally with their neighbours and the overall correlation peaking.

**Figure 10:**
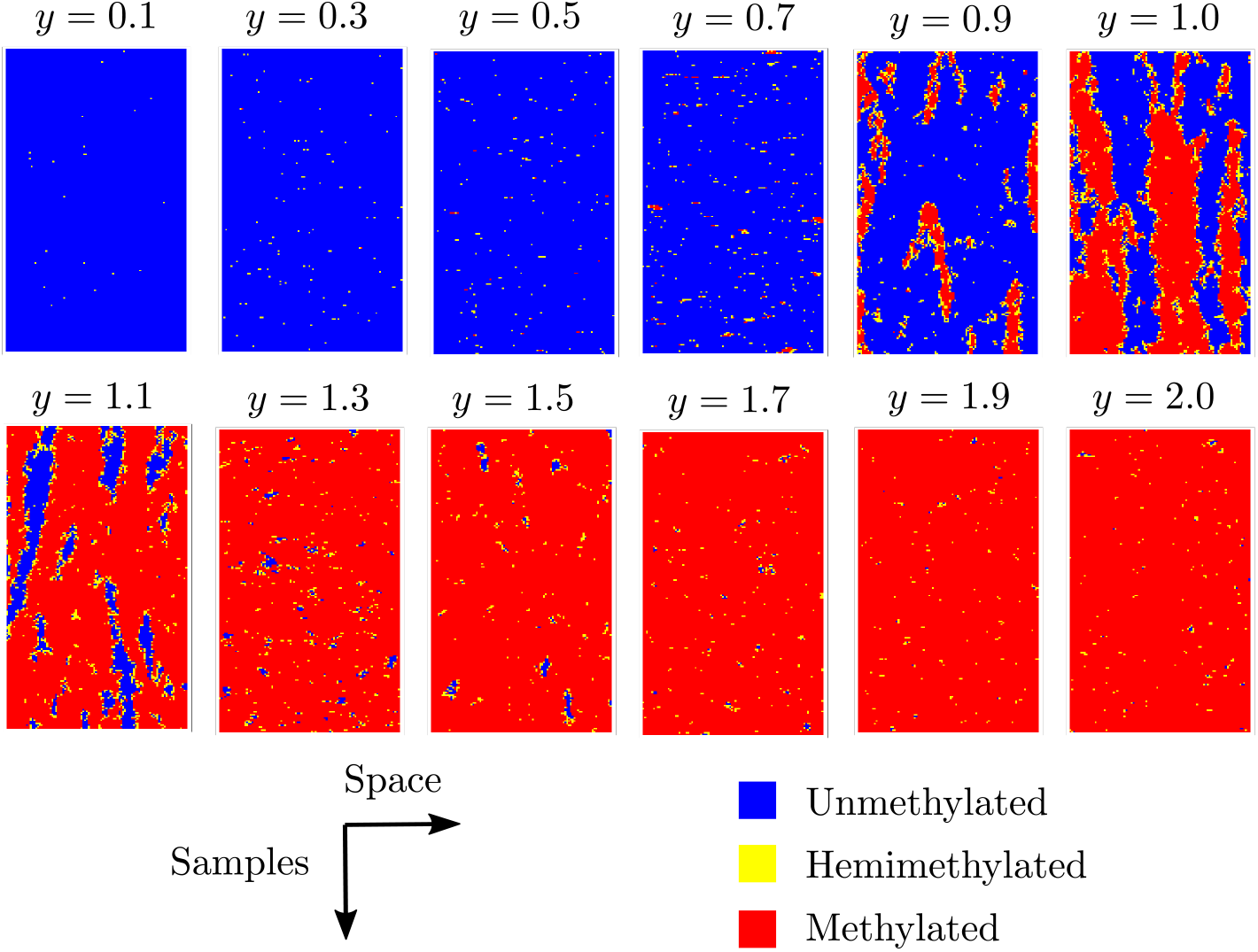
Size of unmethylated (*u*) clusters and methylated (*m*) clusters are most similar when *y* = 1. Simulated methylation patterns are shown for 100 CpGs when *x* = 50. For each *y*, 200 steady-state patterns are displayed, with each row showing the 100 sites at a different time point.

We next consider predictions of these statistics from the MF models. Predictions from the OPMF model lie within the error observed in the simulated data for all parameters considered (Fig. 9). Conversely, predicted statistics from the DPMF model show large deviations from the corresponding simulated statistics when *x* is large and *y* is close to one demonstrating that it has lower predictive power.

### 4.3 Overlapping pairs MF model can infer the parameters underpinning large-scale methylation patterns

To test whether the OPMF model could, in principle, be used to infer collaborativity and methylation strengths from data, we generate synthetic data for selected model parameters, see Section 2.2. We then infer these parameters back using the OPMF model.

Methylomes are typically assayed by whole genome bisulfite sequencing [37]. A variant of this, hairpin-bisulfite sequencing, can be used to assay both strands of each DNA molecule [38]. In both cases, the resulting data is composed of short reads. Each read assays few CpGs and we do not know if reads originate from the same cell or DNA molecule. Simulated datasets from previous sections do not provide a good reflection of bisulfite sequencing since all of the CpGs in the simulated system were sampled at the same time points, the equivalent of the CpGs originating from the same molecule. To obtain short-read data, we instead simulate data for *N* = 1000 CpGs. After steady state is reached, we sample the system at 10, 000 different time points. This is equivalent to sampling 10, 000 molecules in steady state at a single time point. For each CpG, we take the methylation state at 30 time points, randomly chosen from the original 10, 000. The time points chosen for each CpG site are independent of those chosen for other CpGs. This emulates hairpin-bisulfite sequencing data with coverage of 30 reads per CpG. We combine the sample states for all CpGs into a single dataset, *X*, and consider *z* = {*z*_1_, *z*_2_, …, *z*_1000_} where *z*_*i*_ = {1, 2, 3} if *X*_*i*_ corresponds to a *u, h, m* state, respectively. The mean and variance of *z* are calculated and used for inference.

There are numerous well-established methods for conducting inference. For example, inference can be conducted using maximum likelihood estimation [39], which provides point estimates for model parameters. Here we use a Bayesian inference approach, which has the advantage of providing us with a distribution of the estimate value from which we can calculate confidence intervals associated with our inferred parameter values [40, 41, 42, 43]. In particular, we use the Approximate Bayesian Computation Sequential Monte Carlo algorithm (ABC SMC), a likelihood-free inference method (see [44] for a comprehensive overview). We use uniform priors, *U* (0, 100) and *U* (0, 2), for *x* and *y*, respectively. We also define the distance, *d*, between the simulations and model prediction to be the sum of the absolute relative errors of the mean and variance, i.e.

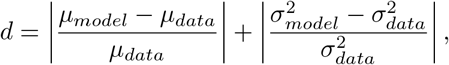

where *µ*_*model*_, *µ*_*data*_ are the means of *z* from the model and data, respectively, and 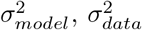, are the variances associated with the model and data, respectively. To rapidly select appropriate tolerances, we calculate the true distances between the simulated data and model predictions at the parameter values of interest.

As in previous sections, we examine *x* = {0.1, 1, 5, 50}. For each *x*, we infer for *y* = {0.3, 1, 1.7} using the GpABC Julia package [45]. Accepted *x, y* values from the final ABC SMC population make up the posterior distributions for *x* and *y*, with the means taken to be the inferred parameter values. 95% confidence intervals were calculated by removing the lowest 2.5% and highest 2.5% from the posteriors.

We find that the inferred parameter values are always of the same order of magnitude as the true values (Table 2), with the most successfully inferred parameters being inferred within 1% of the true values (Fig. 11a, b). There are only two cases where the true parameter values lie outwith the inferred 95% confidence intervals (e.g. Fig. 11c). However, we obtain wide posteriors for large *x* (e.g. Fig 11d), indicating more uncertainty in the inference.

**Table 2:** Inferred parameters for the nearest-neighbour collaborative model using the OPMF model and the ABC SMC algorithm.

| | $y = 0.3$ | $y = 1$ | $y = 1.7$ |
| --- | --- | --- | --- |
| $x = 0.1$ | $(x, y) = (0.197, 0.329)$ | $(x, y) = (0.066, 0.997)$ | $(x, y) = (0.101, 1.701)$ |
| $x = 1$ | $(x, y) = (0.676, 0.251)$ | $(x, y) = (1.011, 0.990)$ | $(x, y) = (0.923, 1.730)$ |
| $x = 5$ | $(x, y) = (6.302, 0.338)$ | $(x, y) = (5.287, 1.000)$ | $(x, y) = (6.432, 1.613)$ |
| $x = 50$ | $(x, y) = (59.571, 0.329)$ | $(x, y) = (50.963, 0.999)$ | $(x, y) = (65.347, 1.599)$ |

**Figure 11:**
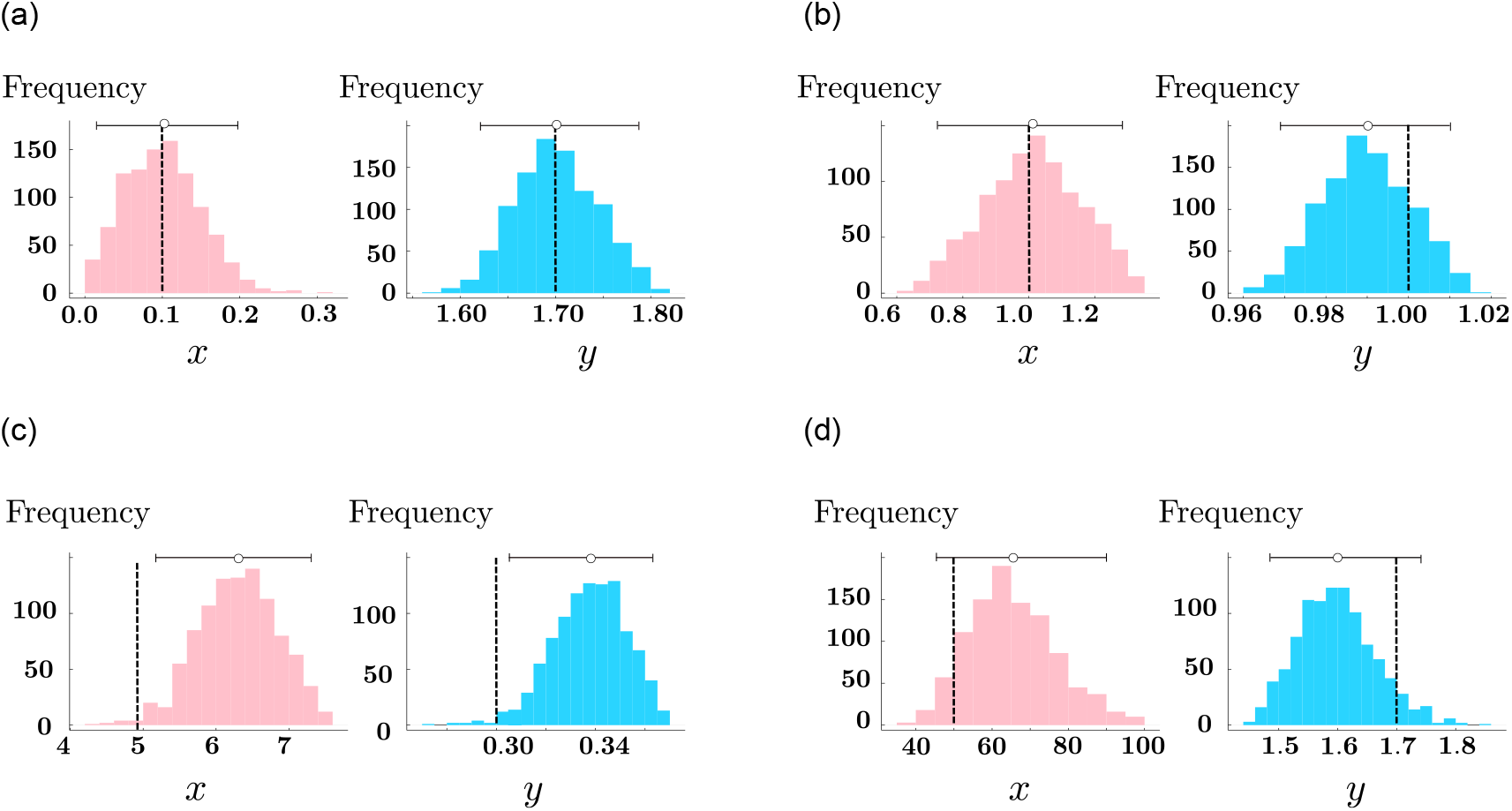
The OPMF model can be used to infer collaborativity and methylation strengths. Example posteriors from inference are shown, with (a), (b), (c) and (d) each corresponding to a different parameter set. True parameter values are denoted by dashed lines, inferred parameter values are shown as dots, with 95% confidence intervals shown as horizontal bars.

The accuracy of inference is highly dependent on the model sensitivity to parameters, with ease of inference increasing as sensitivity to the parameter increases. To test the sensitivity of our model to model parameters, we calculate the relative sensitivity [46] of the OPMF model to the parameters *x* and *y*, for *x* = {0.1, 0.2, … 99.9, 100} and *y* = {0.1, 0.2, … 1.9, 2}. For all parameters considered, the *x*-sensitivity divided by the *y*-sensitivity is strictly less than one, indicating that the model shows more sensitivity to *y* than *x*. This means that *y* will be inferred more accurately than *x*.

## 5 Discussion

Genomic DNA methylation patterns vary between cell types, across differentiation and in disease. The mechanisms underpinning this variation remain unclear but can be better understood using mathematical models. Current approaches are limited by their inability to feasibly model large systems of CpGs and thus understand known large-scale features of methylomes. Here we show that a cluster MF model, based around overlapping pairs of CpGs, can predict DNA methylation patterns generated under a nearest-neighbour collaborative model. This suggests that MF models are a valuable tool for understanding large-scale DNA methylation features.

Previous studies have used mathematical modelling to gain insight into the mechanisms regulating the establishment and maintenance of DNA methylation patterns. In particular, the requirement of collaborativity between CpGs to maintain DNA methylation patterns was postulated through modelling [14] before being observed experimentally [15, 17]. Previous models of DNA methylation rely on stochastic simulations. However, their computational expense limits their use to the study of promoter-scale DNA methylation (regions around 1Kb in size). The stochastic simulations we run here on 200 or 500 sites can take hours to run. In contrast, the OPMF model can be applied to arbitrarily large systems of CpGs and solved numerically in seconds to give accurate predictions for statistics of interest. Our model is based upon the same, or similar, reaction systems used in previous stochastic modelling studies [14, 28, 29], but can be used to study larger systems of CpGs than previously considered. This makes our MF model far better suited to understanding the mechanisms underpinning megabase-sized variations in DNA methylation observed in development, ageing and cancer, which occur at a scale three orders of magnitude larger than promoters [27]. To our knowledge, the largest system previously examined mathematically contained 10^5^ CpGs [15]. However, here simulations were conducted for only a single model parameter set. Running large-scale simulations of this type for many parameter sets will result in computational bottlenecks, meaning that such simulations cannot be used for inference. A previous study has proposed a method, based on the generalized method of moments, for rapid inference using DNA methylation patterns [47]. However, the largest system tackled with this approach contains 10 CpGs. Here we show that our OPMF model can, in principle, be used for accurate, time-efficient inference when modelling arbitrarily large genomic regions.

When used for inference, the OPMF model has a higher sensitivity to methylation strength (*y*) than collaborativity strength (*x*) explaining why the former is generally better inferred than the latter. However, some posteriors obtained in Section 4.3 are very wide and/or the true parameter values lie outside the inferred 95% CIs, indicating that there is scope for inference to be improved. Since our model shows impressive performance in forward prediction, discrepancies between true and inferred parameters are likely due to insufficient data or the inference technique used. However, our analysis demonstrates that the OPMF model can in principle be used for inference on DNA methylation data. It’s accuracy may be improved in future comprehensive studies by experimenting with different inference techniques and sample sizes. In addition, we have inferred parameters using only the mean and variance in methylation state, which can be estimated from standard short-read data. The recent application of long-read technologies to assay DNA methylation patterns [48] could enable the computation of higher order statistics from experimental data, such as the correlation in methylation state between neighbouring sites. Using these additional summary statistics for inference could improve results.

Here we assume that the processes governing the creation of methylation patterns *in vivo* are described well by our nearest-neighbour collaborative model. It is possible that collaborativity *in vivo* can occur between non-nearest-neighbours, something which is not explicitly accounted for in our nearest-neighbour collaborative model. Collaborative methylation interactions are likely determined by the properties of the DNA methylation machinery. DNMT1 and DNMT3B both methylate processively along DNA strands whereas DNMT3A methylates in a distributive manner but can form multimers along the DNA fibre [30]. However, the range and strengths across which these interactions occur is currently unclear so we focus on nearest-neighbour interactions. We note that our OPMF model does capture interactions beyond nearest neighbours because the mean state of pairs in the system influences the change in state of CpGs. Previous modelling studies have also considered different forms of collaborative interactions between CpGs in a system (Fig. 12a,b). In [14] collaboration between any CpGs in the system and nearest-neighbour collaborative methylation alongside distance-dependent collaborative demethylation were both demonstrated to produce stable CpG clusters that were either methylated or unmethylated. Stable clusters were also observed under a distance-dependent collaborative model, where collaborative demethylation dominates over short ranges and collaborative methylation dominates over long ranges [28].

**Figure 12:**
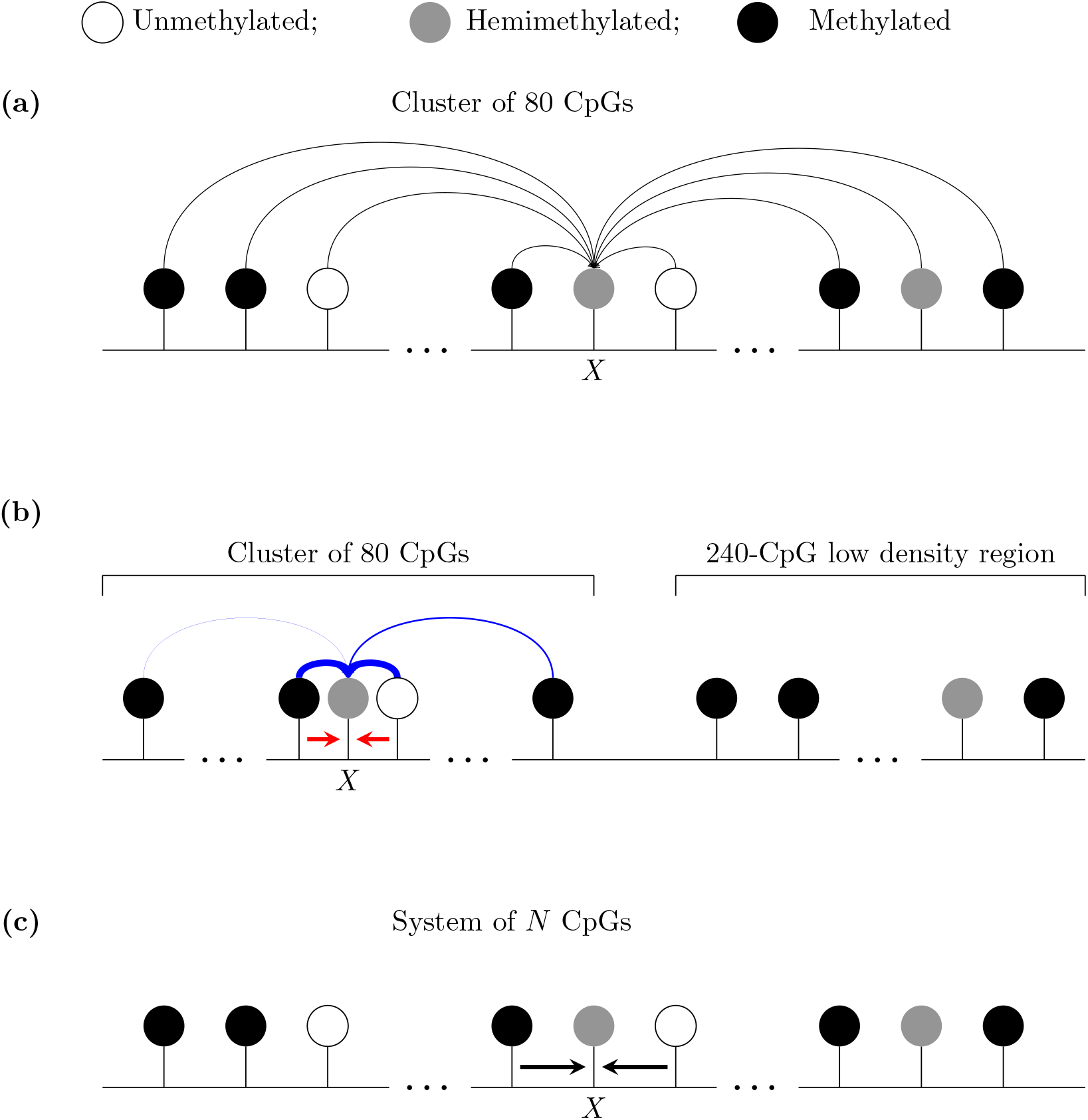
Potential collaborative interactions that can influence a target, *X*, under the models in [14] and the model considered here. (a) A cluster of 80 CpGs is first considered in [14], where a CpG can collaborate with any other CpG in the system with equal probability. (b) A high-density cluster (of 80 CpGs) adjacent to a highly methylated low-density region (of 240 CpGs) is then considered in [14], where there is nearest-neighbour collaborative methylation (red arrows) and the probability of collaborative demethylation occurring due to interaction between two sites decays as the distance between them increases (blue arrows; decay in reaction probability shown by narrowing width of arrows). Note that collaborative demethylation is restricted to the 80-CpG cluster. (c) In this paper collaborative reactions only occur between neighbouring CpGs and there is no upper bound on the system size, allowing large-scale patterns to be considered.

Here we have also assumed that the system of CpGs we model reaches a steady state. This means that we consider either non-dividing cells, or dividing cells which settle down to steady state between replication events such that DNA replication has no long-term effects on DNA methylation patterns. Whether or not a cell satisfies the latter case is dependent on the real magnitudes of *k*_*i*_, *i* = {1, …, 12}, and the time between replication events. Experimental studies show that arrested cells have similar DNA methylation patterns to those that are cycling, supporting the assumption that DNA replication has no long-term effect on DNA methylation [31]. Furthermore, an analysis of DNA methylation patterns on newly synthesised DNA suggests that re-methylation occurs within 20 minutes of replication [49]. However, another analysis of DNA methylation following replication suggests re-methylation is often delayed [50]. At present, it is unclear whether this delay is sufficient to have an effect on methylation patterns during the following cell cycle.

Our assumption that *k*_*i*_, *i* = {1, …, 12} take the form in Table 1 could be violated in reality. For example, DNMT1 shows a strong preference for *h* sites over *u* sites [30], meaning that methylation reactions with an *h* target may have higher reaction rates than those with a *u* target. Future work could relax rate assumptions to account for such factors. Preliminary investigations confirm that the OPMF model provides a good approximation to the nearest-neighbour collaborative system when *k*_*i*_, *i* = {1, …, 12} are considered as twelve independent parameters (data not shown). However, the difficulty of parameter inference increases with the number of parameters, meaning that relaxing parameter assumptions will likely decrease inference quality. Nonetheless, our model suggests that it is the parameters *x* and *y* that determine steady-state methylation patterns, rather than individual reaction rates. This is supported by a study where modelling of experimental data suggested that the ratio between methylation and demethylation rates determines steady-state methylation levels at single CpGs [51].

Here, we demonstrate that MF models can accurately predict the behaviour of large CpG systems subjected to nearest-neighbour collaboration. Our study presents the first mathematical modelling approach that can be applied to arbitrarily large systems of CpGs. The future application of this approach will facilitate the delineation of the methylation dynamics that underpin the formation of large-scale methylation patterns in developmental and disease contexts.

## 6 Acknowledgements

We thank Chris Ponting, Diego Oyarzun and members of the Grima and Sproul lab for helpful discussions about the manuscript. LK is a cross-disciplinary post-doctoral fellow supported by funding from the University of Edinburgh and Medical Research Council (MC UU 00009/2). DS is a Cancer Research UK Career Development fellow (reference C47648/A20837), and work in his laboratory is also supported by an MRC university grant to the MRC Human Genetics Unit. RG is supported by Leverhulme Trust research awards (RPG-2018-423 and RPG-2020-327).

## 7 Contributions

LK conducted the modelling and analyses presented in the manuscript. DS and RG planned and supervised the study. LK, DS and RG wrote the manuscript.

## Appendix

### A Derivation of effective reaction rates for the DPMF model

Here we illustrate how *â*_1_, the reaction rate associated with *mm hm* → in (18), is constructed. We first consider all reactions in Fig. 1 that can occur *within* the *mm* pair, resulting in conversion to an *hm* pair. Clearly, *â*_1_ must contain a 2*k*_3_ term since *mm* → *hm* occurs if either *m* undergoes 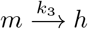. Note that *mm* → *hm* cannot occur due to a collaborative interaction between the two *m* sites in *mm*.

Next, we consider all *mm* → *hm* reactions that can occur due to an *m* within the *mm* interacting with a site from an adjacent pair. For example, an *m* could collaborate with a *u* from an adjacent pair via 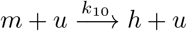, i.e. we can have

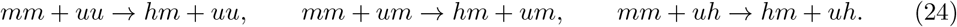

The first reaction can be written as an effective first-order reaction, 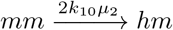, where the *uu* is absorbed into the reaction rate by making it proportional to *µ*_2_, and the factor of two allows for either *m* within the *mm* to undergo this reaction. The second reaction in (24) can be written as 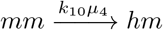, where the *um* is absorbed into the reaction rate and either *m* can undergo the reaction, giving us a factor of two. However, this reaction requires the *u* within the *um* to be directly adjacent to the *mm*. The probability of having *um* in this particular order is 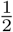, giving an effective reaction rate of 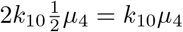. Similarly, the third reaction in (24) can be written as 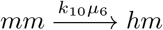.

Finally, an *m* within the *mm* can interact with an *h* from an adjacent pair, via 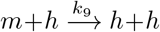i.e. we can have

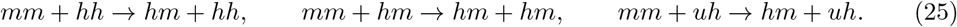

Using similar arguments as above, we can write each of these three reactions as the effective first-order reaction *mm* → *hm*, with rates 2*k*_9_*µ*_3_, *k*_9_*µ*_5_ and *k*_9_*µ*_6_, respectively.

Hence,

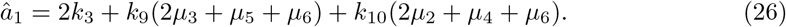

### B Derivation of effective reaction rates for the OPMF model

We now construct *ã*_1_, the reaction rate associated with *mm* → *hm* in (23). As with the DPMF model, reactions occurring within the *mm* pair contribute a 2*k*_3_ term to *ã*_1_.

Now, *mm* → *hm* can occur due to a 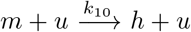 reaction if one of the *m* sites in the *mm* pair is also in a pair with a *u* site, i.e. we can have

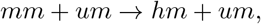

where now the *um* and *mm* reactants share a common *m*. This can be written as the effective first-order reaction *mm* → *hm* with rate

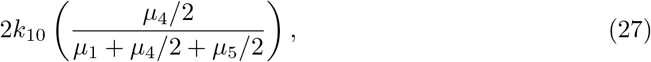

Where 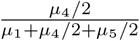 is the probability that an *m* within the *mm* also forms a *um* with its other neighbour, i.e. it is the conditional probability that a pair is in the *um* state given that we know a particular site in the pair is *m*. Recall that *µ*_4_ gives the proportion of *um* and *mu* pairs, while *µ*_5_ gives the proportion of *hm* and *mh* pairs. The factors of 1*/*2 in (27) account for the fact that the *u* and *h* in the *um* and *hm* must be on a particular side of the *m* (since an *m* is on its other side). The factor of two at the front of (27) allows for either *m* in the *mm* to undergo the reaction.

Similarly, *mm* → *hm* can occur due to a 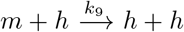 reaction if one of the *m* sites in the *mm* is also in a pair with a *h* site, i.e. we can have

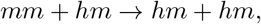

where the reactant *hm* and *mm* share a common *m*. This reaction can again be written as *mm* → *hm*, where the rate,

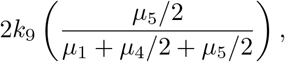

is derived in a similar way as above. Hence, we have

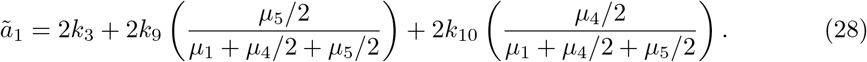

### C Moment equations for the cluster MF models

Following the approach in Ref [33] (see Appendix C) we obtain first moment equations for the DPMF and OPMF models. Here *µ*_1_—*µ*_6_ are the mean levels of each paired state. Note that, for *i* = 1, …, 12, *a*_*i*_ = *â*_*i*_ in the DPMF model and *a*_*i*_ = *ã*_*i*_ in the OPMF model,

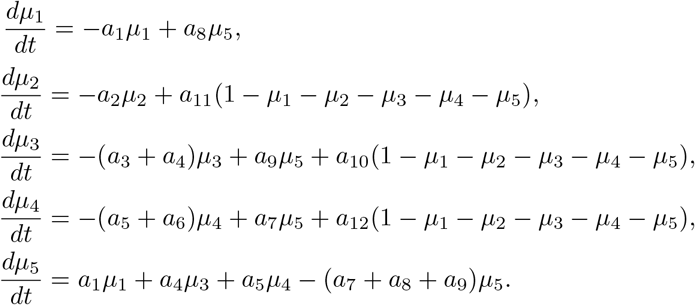

Note again that we can obtain *µ*_6_ via *µ*_6_ = 1 − *µ*_1_ − *µ*_2_ − *µ*_3_ − *µ*_4_ − *µ*_5_. The second moment equations, describing the evolution of ⟨*L*_*i*_*L*_*j*_⟩ for 1 ≤ *i, j* ≤ 5, are given by

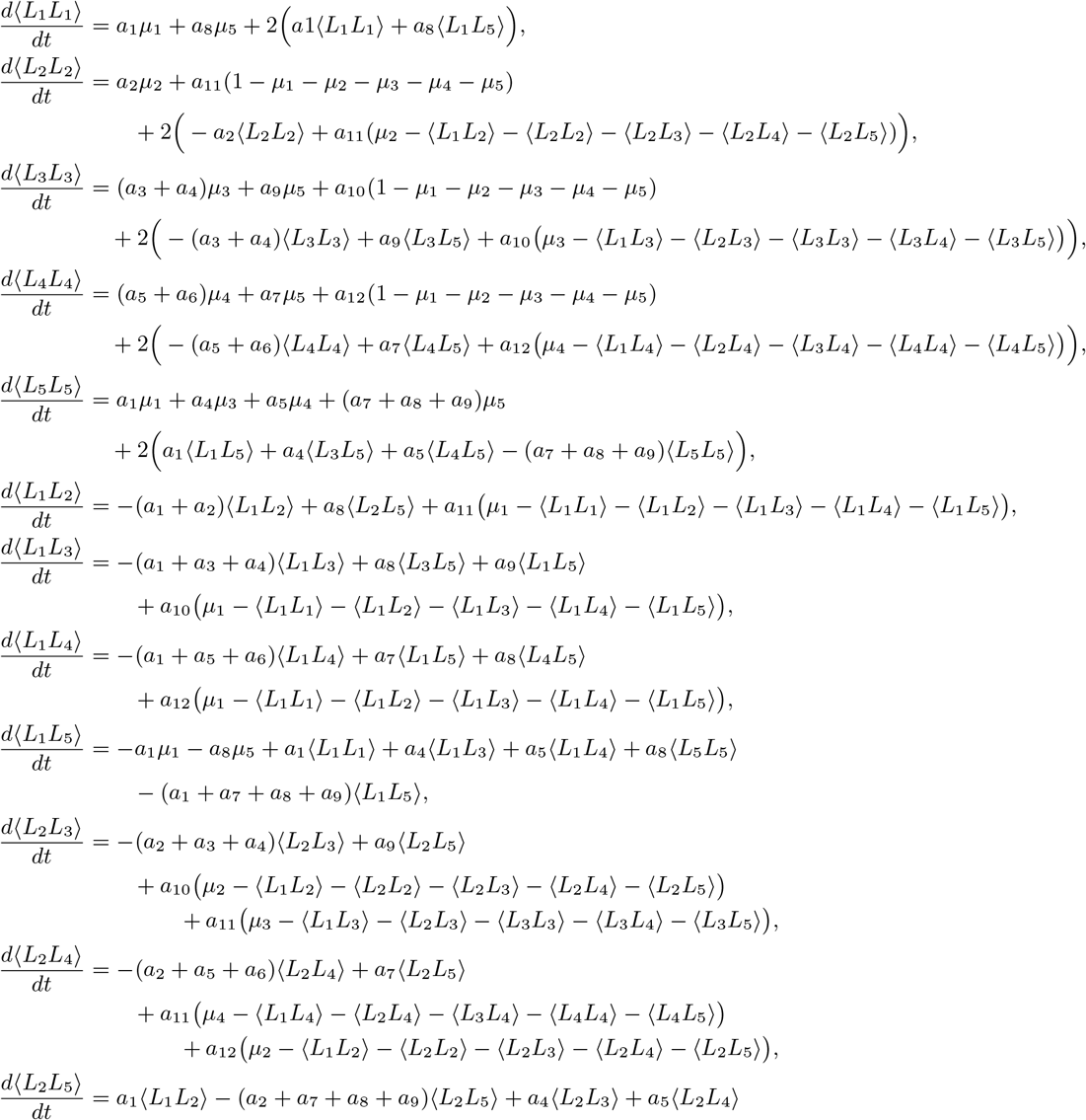

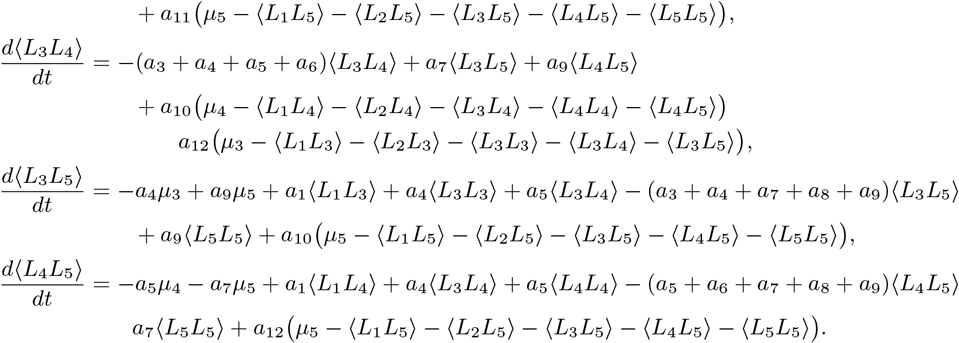

Using *L*_6_ = 1 − *L*_1_ − *L*_2_ − *L*_3_ − *L*_4_ − *L*_5_, we obtain

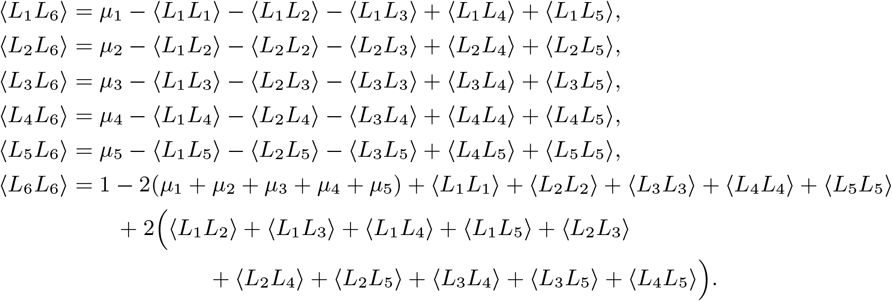

